# Objective but not subjective fatigue increases cognitive task avoidance

**DOI:** 10.1101/208322

**Authors:** Charles-Etienne Benoit, Oleg Solopchuk, Guillermo Borragán, Alice Carbonnelle, Sophie Van Durme, Alexandre Zénon

## Abstract

Mentally demanding tasks feel effortful and are usually avoided. Furthermore, prolonged cognitive engagement leads to mental fatigue, consisting of subjective feeling of exhaustion and decline in performance. Despite the intuitive characterization of fatigue as an increase in subjective effort perception, the effect of fatigue on effort cost has never been tested experimentally. To this end, sixty participants in 2 separate experiments underwent a forced-choice working memory task following either a fatigue-inducing (i.e. Stroop task) or a control manipulation. We measured subjective fatigue and effort as well as their objective behavioral signatures: performance decline and task avoidance, respectively. We found that fatigue-induced performance decline was correlated with task avoidance, while the feelings of fatigue and effort were unrelated to each other. Our findings highlight the discrepancy between subjective and objective manifestations of fatigue and effort, and provide valuable evidence feeding the ongoing theoretical debate on the nature of these constructs.

## 1. Introduction

Demanding cognitive activities are typically aversive and lead to the percept of effort (Inzlicht et al., 2015; Westbrook and Braver, 2015). The subjective, introspective dimension of cognitive effort is assessed by means of self-report questionnaires, such as the NASA task-load index (Hart and Staveland, 1988), but cognitive effort can also be evaluated more objectively by assessing task avoidance. This can be done by asking participants to choose between alternative options involving tasks of varied difficulty, associated with rewards of varied magnitude and by using neuroeconomic models to evaluate the effort cost of each task, i.e. how much performing the task discounts the associated reward (Hosking et al., 2014; Westbrook et al., 2013).

While the existence of a cost to cognition appears intuitively obvious, its origin remains unknown (Kurzban et al., 2013; Shenhav et al., 2017; Westbrook and Braver, 2015). What is clear, however, is that prolonged cognitive engagement leads to the subjective feeling of fatigue (Ackerman and Kanfer, 2009; Campagne et al., 2004; Deluca, 2005) and can also deteriorate performance (Holtzer et al., 2011; Schwid et al., 2003; Tanaka, 2015), sometimes referred to as objective fatigue (Bailey et al., 2007). The nature of these behavioral manifestations of cognitive fatigue is diverse, including disruption of inhibition mechanisms (Kato et al., 2009), planning (Lorist, 2008; Lorist et al., 2000; van der Linden et al., 2003), processing of new information (Massar et al., 2010), attention (Boksem et al., 2006; Dorrian et al., 2007; Holtzer et al., 2011; Lim et al., 2010), working memory (Gergelyfi et al., 2015) or task switching (Borragán et al., 2017). Like cognitive effort, the origin of cognitive fatigue remains unclear. Cognitive fatigue is hypothesized to result from metabolic depletion (Christie and Schrater, 2015), spreading of local sleep patterns in the brain (Krueger et al., 2008; Vyazovskiy et al., 2011), the buildup of a decision signal aimed at limiting the opportunity cost of engaging cognitive resources in the same task (Kurzban et al., 2013) or by the extended use of high-consuming strategies aimed at maintaining performance (i.e. Compensatory Control Model (CCM); Hockey, 2013, 1997).

One puzzling and pervasive observation in the fatigue literature is that the subjective dimension typically fails to correlate with the behavioral consequences of fatigue (Deluca, 2005). According to Hockey’s influential CCM model (Hockey, 2013, 1997), this lack of relation between the subjective and objective dimensions would be explained by the fact that subjective fatigue is the manifestation of compensatory resource mobilization, needed to maintain performance despite the progressive deterioration of capacity induced by fatigue. Therefore, initially, subjective fatigue would predominate, while objective performance decrement would kick in later on in the building up of fatigue. Subjective feelings of fatigue would appear as an indication that the resources needed to accomplish the ongoing demands are at risk or no longer available. Neurophysiological evidence for increased engagement during the compensatory phase of cognitive fatigue has been recently reported (Wang et al., 2016).

Phenomenologically, subjective fatigue is often reported as increased aversion for cognition and increased percept of effort (Arai, 1912). In addition, the CCM assumes that subjective fatigue should be accompanied with increased percept of cognitive effort caused by the compensatory engagement of resources. However, to the extent of our knowledge, the link between fatigue and effort, in both their subjective and objective dimensions, has never been experimentally assessed. We hypothesized that subjective fatigue would increase the cost of effort: a fatigued person would feel more effort performing a task of equivalent difficulty and would increase their preference for low-demand tasks. Sixty participants engaged in 2 separate experiments in which subjective effort and task avoidance were measured following either fatigue-inducing or control manipulation. We evaluated fatigue both in its subjective dimension and in its manifestation as performance decline, and controlled for other variables such as sleepiness, arousal and motivation. We found that objective fatigue increased task avoidance, while subjective feelings of fatigue and effort were unrelated to each other.

## 2. Experiment 1

### 2.1. Materials and methods

#### 2.1.1. Participants

Thirty right-handed healthy participants took part in Experiment 1 (27 F, 3 M, Age: 22.8 +- 2.5, mean, SD), receiving monetary compensation for their participation (70 – 100€). Each experiment was performed over the span of three consecutive days. This included training on day 1 and fatigue/control manipulation followed by the working memory task on the experimental days 2 and 3, all occurring at the same time during the day to control for circadian effects between sessions. All participants reported a normal or corrected-to-normal vision, had no auditory impairments or neurological disorders. Experiments were carried out according to the Declaration of Helsinki and were approved by the Ethics Committee of the Université catholique de Louvain. All participants provided a written informed consent.

#### 2.1.2. Subjective evaluation

Participant’s fatigue was assessed using the French version of the multidimensional fatigue inventory (MFI) (Gentile et al., 2003), composed of fifteen statements to be rated on a 5-point Likert scale (from “1 - Yes, that is true” to “5 - No, that is not true”). The answers to all questions were summed to obtain a Fatigue Index, in which lower score reflected higher fatigue. MFI was evaluated at the beginning and at the end of both experimental days.

In order to assess sleepiness, we used the 9-point Karolinska sleepiness scale (KSS) (Åkerstedt et al., 2014; Åkerstedt and Gillberg, 1990). Participants had to select a number corresponding to their level of sleepiness (from 1 = extremely alert to 9 = very sleepy, fighting against sleep). KSS was evaluated simultaneously with the MFI.

We also evaluated participants’ intrinsic motivational state using intrinsic motivation inventory (IMI, http//:selfdeterminationtheory.org/intrinsic-motivation-inventory/) The subjects provided the ratings on a 7-point Likert scale (from “Not at all true” to “Very true”), thus, higher scores represented higher task interest/enjoyment.

Subjective effort was evaluated using the National Aeronautics and Space Administration Task Load Index (NASA-TLX) rating scale (Hart and Staveland, 1988). It is divided in six subscales: mental demand, physical demand, temporal demand, performance, effort and frustration level. In the present study, we focused only on mental demand, performance and effort. Since the concepts of mental demand and effort were difficult to tease apart for participants, and since the scores strongly correlated with each other, we took the average of the mental demand and effort scores as an index of subjective effort cost. The participants had to score each of the items on a 20-point scale presented directly on screen. Participants completed the NASA-TLX after each block of the working memory task on both experimental days.

#### 2.1.3. Fatigue/Control manipulation

On experimental days, participants underwent 2h of either fatigue or control manipulation (see Figure 1). Fatigue was induced by performing a variant of the Stroop task (MacLeod, 1991), widely used as a way of promoting cognitive fatigue (Moeller et al., 2012; Pageaux et al., 2015; Wang et al., 2016). In the control session, participants watched emotionally neutral documentaries. The order of the days was counterbalanced between all participants. In the Stroop task, participants were instructed to be as accurate and fast as possible. The time between the cue onset and the initiation of the next trial was adapted to the performance of each participant, according to the following rule:

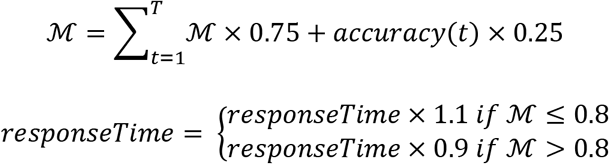

with *T* being the current trial number, *responseTime* being initialized to 1500 ms at the beginning of each block, 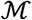 being intialized to zero, and *accuracy(t)* representing the accuracy of the response in trial *t* (one if correct, zero if incorrect). `responseTime` was 300 ms minimum and 3000 ms maximum. This procedure ensured that the task remained challenging for all subjects, independently of their capacity. In our modified version of the Stroop task, each block contained a total of 368 trials, distributed equally between 2 different tasks. In the number task, participants were presented with two numbers, on either side of the fixation point (left and right), and had subsequently to report the location of either the largest in value or the largest in size, depending on an instruction cue. In the arrow task, one arrow was presented either above or below the fixation point and participants were to report either its position or direction, depending on the instruction cue. Trials were equally divided into congruent (e.g. large digit with large value or upwards pointing arrow located on the upper half of the screen) and incongruent (e.g. large digit with small value or upwards pointing arrow located on the lower half of the screen) trials. The participants completed 8 blocks in 2 hours. Each trial proceeded as follows: the fixation cross appeared in the middle of the screen for 1000 ms, followed by 250 ms of stimuli presentation. After a delay, a blank screen appeared for 500-1000 ms (depending on the participant performance), and the instruction cue (“arrow” task: “direction” or “location”; “number” task: “value” or “direction”) was presented for 550 ms. Participants had to answer within the cue presentation time, equal to the *responseTime* variable detailed above. The next trial began right after participants' response.

**Figure 1.**
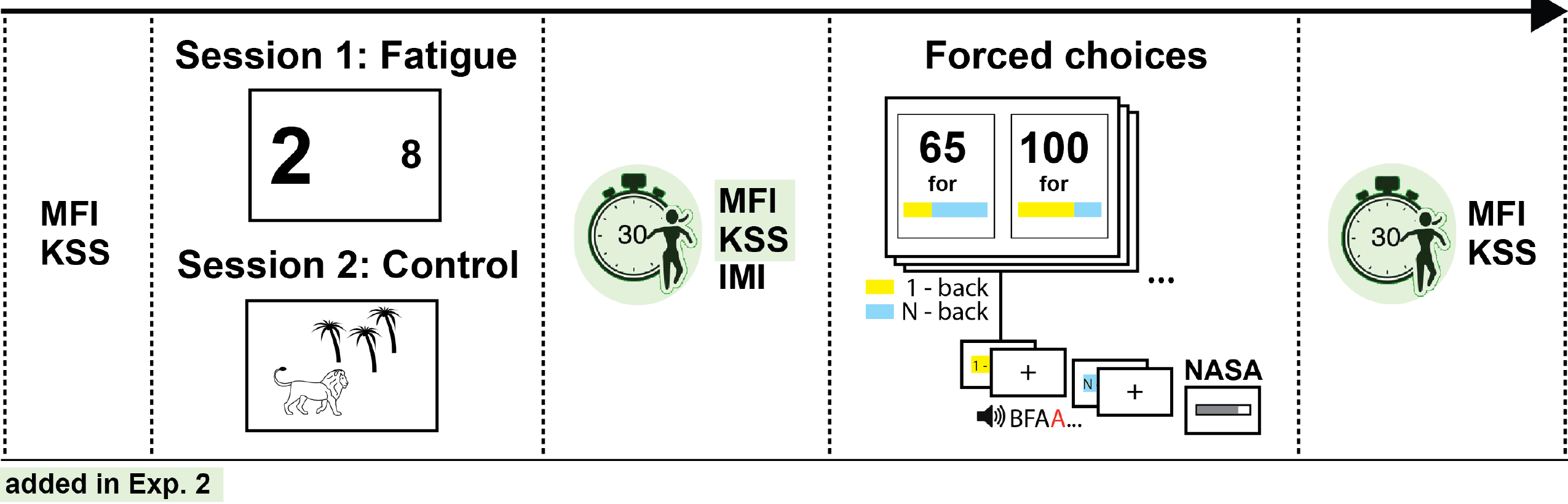
Schematic description of the protocol. Participants were first evaluated with the multidimensional fatigue inventory (MFI) and Karolinska sleepiness scale (KSS). Following fatigue-inducing or control manipulation they again completed the questionnaires (only intrinsic motivation inventory (IMI) in Exp 1), and performed a forced-choice working memory task followed by the National Aeronautics and Space Administration Task Load Index (NASA-TLX). Experiment 2 also included arousal boosts consisting of 30-second run on the spot, prior to questionnaires completion.

During the control session, participants watched animal documentaries on a computer for 2h. The documentaries were chosen according to their preference out of 7 different movies. Similar approaches have already been used in other experiments as a control condition (Badin et al., 2016; Van Cutsem et al., 2017; Zering et al., 2017).

#### 2.1.4. Effort discounting evaluation

All tasks were implemented using version 3.0.9 of the Psychotoolbox (Brainard, 1997) in Matlab 7.5 (The MathWorks, Inc. Natick, Massachusetts, United States). Participants faced a 19-inch CRT screen with a refresh rate of 100Hz. The distance between the screen and the chin support was 58 cm.

To evaluate effort and assess the behavioral consequences of fatigue, we used a variant of the auditory “N-back” working memory task, in which the subjects are required to judge whether the presented letter matches the one heard “N” letters before (Kirchner, 1958). Series of letters from the French alphabet, excluding F, J, M, P, W, Y, were presented aurally to participants. In each block, 25% of the letters were defined as targets, the others being considered as distractors. Task difficulty ‘N’ was calibrated individually during the training on the first day, and stayed constant during the whole experiment. The training included 15 blocks of increasing demand (N = 1 to 5), composed of 60 letters each. The letters were presented continuously while participants fixated the cross in the center of the screen. Response timeout was fixed to 1.5 seconds (indicated by change of fixation cross orientation), followed by an inter-stimulus interval of 2 to 3 seconds (uniformly distributed). A short pause was provided after every 5 blocks. The answers were collected via computer keyboard (distractor and target corresponded to button number ‘1’ and ‘2’ respectively). When the participant performance was greater than or equal to 70% (mean of the % of targets detected and the % of distractors correctly rejected) they progressed to the next level with a maximum difficulty level set to 5-back, otherwise they repeated the same level once more.

Following the fatigue or control manipulation on experimental days, participants performed the N-back tasks of different difficulty levels, presented in the context of a forced-choice paradigm, inspired from earlier studies (Westbrook et al., 2013). In order to avoid that participants’ preference for difficulty-reward combinations would be influenced by their auto-evaluation of performance, rather than by effort perception, we defined task difficulty as the ratio of 1-back and N-back tasks in the block of 80 letters (actual ‘N’ depending on participants' performance during training, cfr. supra). These ratios are referred to as the block difficulty level (DL): % of N-back: DL-1 = 25%, DL-2 = 37.5%, DL-3 = 50%, DL-4 = 62.5%, DL-5 = 75%). The difficulty was represented visually using horizontal color bars (blue = 1-back; orange = n-back; see Figure 1). A fixed amount of 2 € was always associated with the DL-5 block while a variable amount, ranging between 0 and 2 € (with 0.20 € intervals) was offered for the easy option (i.e. DL-1 to DL-4). Participants had to make the choice by pressing the right or left mouse button depending on whether they wanted to execute the task with the DL presented on the right or left part of the screen. The side of easy and difficult offers was pseudo-randomized for every participant. All possible permutations were evaluated, which resulted in 44 choices. Within these selections, participant had to perform 15 of the randomly selected choices. In each block, the accuracy obtained in both levels (1-back and N -back) and conditions (target hits/distractors correct rejections) was averaged to calculate the performance. This particular procedure ensured that, regardless of DL, performance in the 1-back and in the N-back tasks had the same impact on the amount of money earned at the end of the block. This provided incentives for participants to disregard their performance, and to consider only cognitive effort associated with the DL when making their choice. The remuneration was calculated by multiplying the performance by the reward attributed to the selected block (i.e. 0 to 2 €). This information was provided on the screen at the end of each block. Prior to receiving performance feedback, participants had to answer the NASA-LTX questionnaire, presented on the computer screen.

#### 2.1.5. Statistical analyses

We analyzed data with Matlab (The MathWorks, Inc., Natick, MA, USA) and performed statistical analyses with JASP (Version 0.8.2, JASP Team 2017). We performed both frequentist and Bayesian repeated-measure ANOVAs and Kendall correlations, to avoid undue influence of outliers. Results from the frequentist analyses are reported in terms of p-value (and F statistics for the ANOVA), whereas Bayesian analyses resulted in Bayes Factors (BF). BF represents the ratio of the likelihood of two different models with, most classically, one assuming the alternative hypothesis and the other assuming the null hypothesis. The BF provides a better estimate of effect reliability than the p-value. In addition, it provides the ability to evaluate the probability that the null hypothesis is true. The BF can be interpreted on the basis of the table proposed in Kass and Raftery (1995):

| $\log BF_{10}$ | $BF_{10}$ | Evidence against $H_0$ |
| --- | --- | --- |
| 0 to 1/2 | 1 to 3,2 | Not worth more than a bare mention |
| 1/2 to 1 | 3,2 to 10 | Substantial |
| 1 to 2 | 10 to 100 | Strong |
| > 2 | > 100 | Decisive |

For 1-way ANOVAs and correlation analyses, the BF_10_ is reported, which represents the comparison between the alternative and the null hypothesis. For 2-way ANOVAs, BF_inclusion_ is reported instead. BF_inclusion_ is obtained by comparing the likelihood of the model including, or excluding a given effect.

The behavior of the participant in the choice task was analyzed with a computational model of effort-based decision making inspired from earlier studies (Zénon et al., 2016)

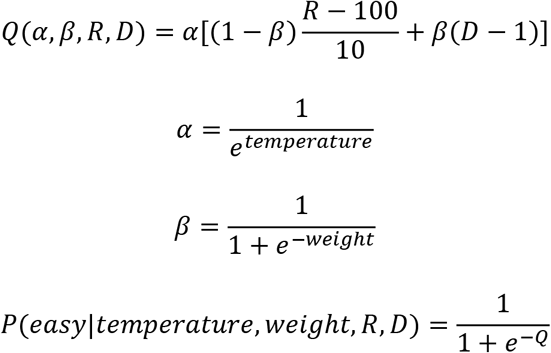

with *temperature* and *weight* as free model parameters. *R* represented the amount of reward proposed for the difficult version of the task, and *D* accounted for the difficulty level. *P(easy)* indicated the probability of choosing the easy option, which was a function of the temperature and weight parameters and of the reward and difficulty conditions. The model was fit with multidimensional nonlinear minimization (Nelder-Mead, fminsearch) with ridge regularization.

### 2.2. Results

#### Stroop session increases MFI and subjective effort but not task avoidance or performance

We performed a one-way repeated-measure ANOVA on the post-pre difference in MFI, KSS and IMI scores with SESSION as factor (control vs fatigue). There was no effect of SESSION on either KSS (F(1,29)=2.188, p=0.15, BF_10_=0.759) or IMI (F(1,29)=1.965, p=0.172, BF_10_=0.672) and a small influence of SESSION on MFI (see Figure 3A; F(1,29)=5.075, p=0.032, BF_10_=1.893; i.e. significant according to frequentist statistics but anecdotal for Bayesian statistics). However, there was a very strong positive correlation between fatigue-control changes in MFI and KSS (Kendall's tau=0.508, p<0.0001, BF_10_=283.292), suggesting that the subjects failed to distinguish clearly the subjective feeling of fatigue from that of sleepiness. Finally, neither KSS nor MFI correlated with IMI (all p > 0.555, all BF_10_ < 2.33), indicating that subjects’ intrinsic motivation was not affected by fatigue manipulation, in accordance with previous studies (Gergelyfi et al., 2015).

The NASA effort and performance scores changed as a function of the DIFFICULTY level (2-way RM-ANOVA: NASA effort: F(4,116)=9.564, p<0.001, BF_inclusion_=4281, NASA performance: F(4,116)=4.767, p=0.001, BF_inclusion_=17.9). In addition, NASA effort, but not NASA performance changed with SESSION (see Figure 3C; NASA effort: F(1,29)= 0.963, p=0.041, BF_inclusion_=361, NASA performance: F(1,29)=.963, p=0.335, BF_inclusion_=0.228). There was no SESSION × DIFFICULTY interaction (NASA effort: F(4,116)=0.180, p=0.949, BF_inclusion_=0.025, NASA performance: F(4,116)=0.163, p=0.956, BF_inclusion_=0.027). There was no correlation between the fatigue-related changes in the two variables (Kendall's tau=-0.080, p=0.547, BF_10_=0.284).

The temperature and weight parameters from the task avoidance models were also subjected to a one-way repeated-measure ANOVA with SESSION as factor (control vs fatigue). The weight parameter can be interpreted as an index of effort cost, while temperature indicates the level of randomness of participants’ behavior. There was no change in either parameters between sessions (see Figure 2, left panel, and Figure 3B; temperature: F(1,29)=0.179, p=0.676, BF_10_=0.285, weight: F(1,29)=0.677, p=0.418, BF_10_=0.353). There was a strong correlation between the fatigue-control changes in the two variables (Kendall's tau=0.526, p<0.001, BF_10_=704.0). However, we noticed that many participants systematically picked the difficult task, irrespective of the reward value (13 subjects picked the difficult task in 100% of choices in at least one session). On average the percentage of choice of the difficult task was 86.4±2.7% and 87.4±2.6% for the fatigue and the control sessions, respectively (mean±CI).

**Figure 2.**
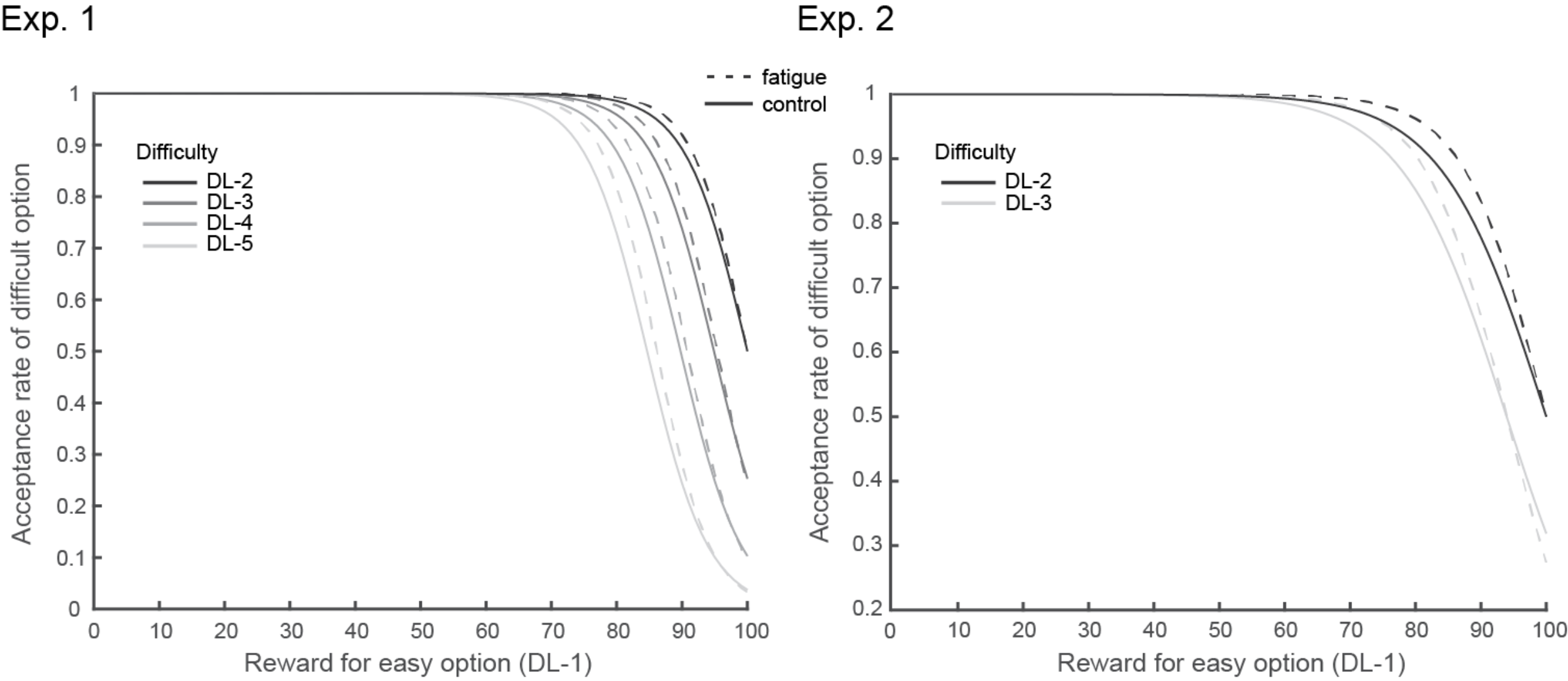
Fitting curves from the behavioral model, averaged across participants for each difficulty level and session. Task preferences indicate a clear effect of difficulty level on probability of accepting more difficult task, with no clear difference between fatigue and control sessions in both experiments.

A 2-way RM-ANOVA on the logit-transformed accuracy scores showed, besides the obvious difference between 1-back and N-back TASK (accuracy: F(1,29)=165.363, p<0.001, BF_10_>10^29^; RT: F(1,29)=83.510, p<0.001, BF_10_>10^16^), an absence of main SESSION effect (accuracy: F(1,29)=2.475, p=0.127, BF_10_=0.374, RT: F(1,29)=1.124, p=0.298, BF_10_=0.354) or interaction between the TASK and the SESSION effect (accuracy: F(1,29)= 2.651, p=0.114, BF_10_=0.550; RT: F(1,29)=0.167, p=0.686, BF_10_=0.266). There was a moderate correlation between the fatigue-control changes in accuracy and RT (Kendall's tau=0.360, p=0.007, BF_10_=7.747).

#### Changes in performance correlate with changes in task avoidance

We found strong evidence for a correlation between fatigue-induced changes in Model Weight and performance in the N-back task (see Figure 4A; Tau=0.416, p=0.001, BF_10_=35.265) but not with 1-back performance (see Figure 4A; Tau=0.190, p=0.162, BF_10_=0.643). In contrast, Model Weight failed to correlate with MFI (Tau=0.009, p=0.943, BF_10_=0.236), KSS (Tau=-0.054, p=0.691, BF_10_=0.256) or NASA scores (see Figure 4B; effort: Tau=-0.117, p=0.376, BF_10_=0.351; performance: Tau=0.218, p=0.094, BF_10_=0.938) changes. There was no correlation between fatigue-induced changes in performance in the 1-back or N-back tasks and NASA performance or effort scores (see Figure 4C; 0.2<BF_10_<0.3).

**Figure 3.**
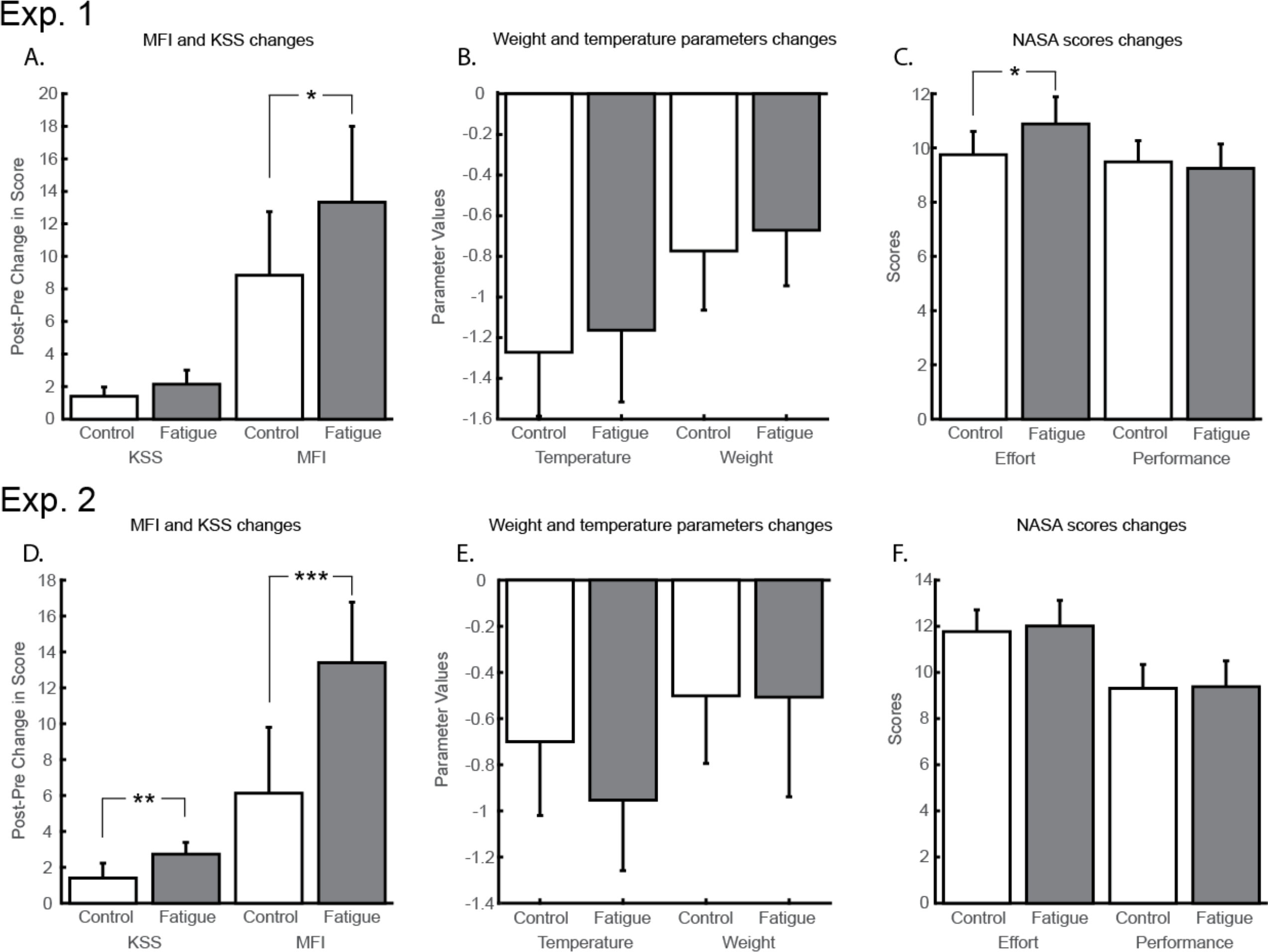
Subjective evaluations and free model parameters in fatigue (gray bars) and control conditions (white bars) for Experiment 1 (A-C) and 2 (D-F). Error bars indicate the 95 % confidence interval. *p < .05. **p < .01.***p < .001.

**Figure 4.**
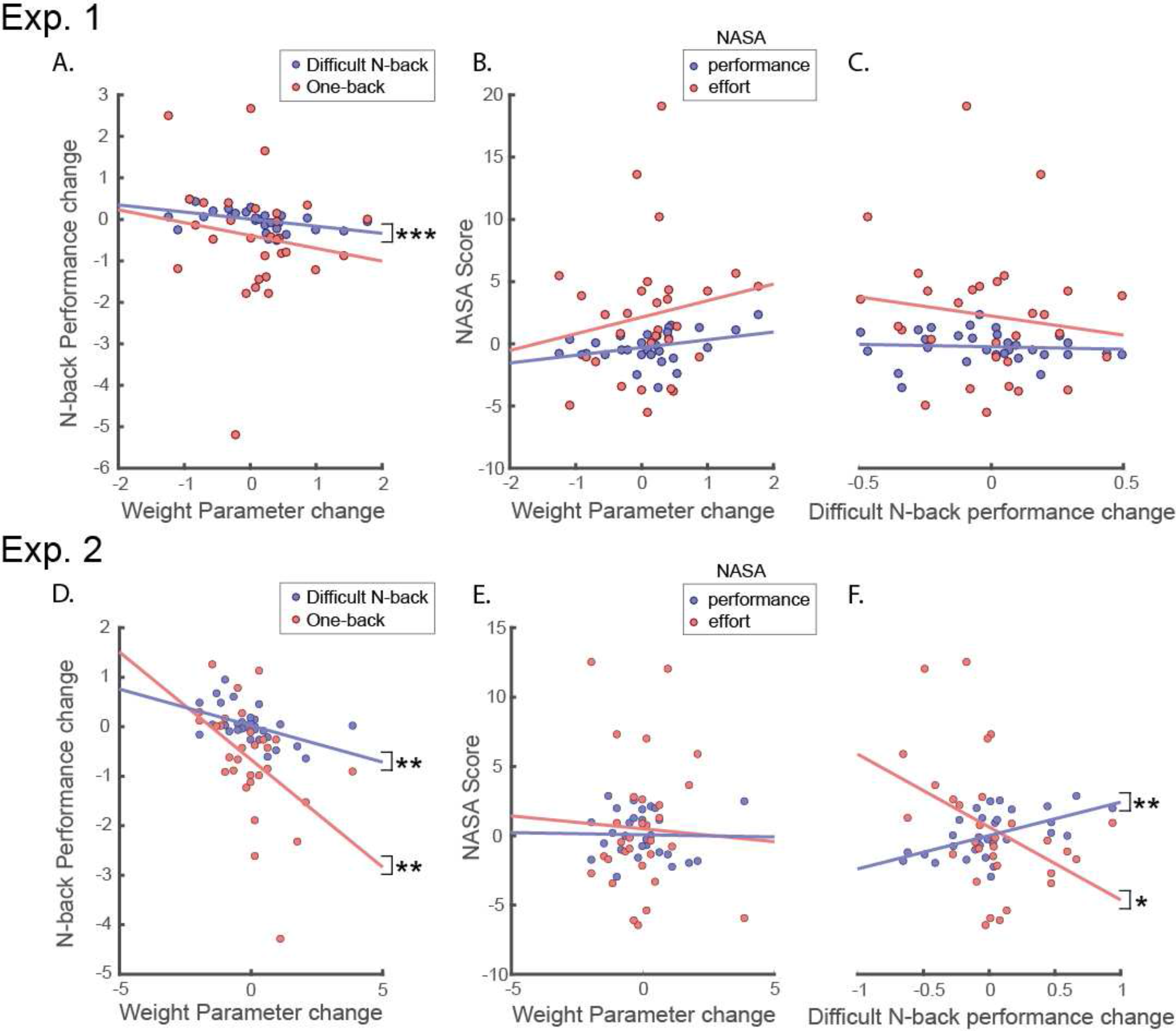
Relationships between session changes in N-back performance, subjective evaluations and free model parameters for Experiment 1 (A-C) and Experiment 2 (D-F). *p < .05, **p < .01, ***p < .001.

#### Subjective fatigue correlates with changes in subjective effort and sleepiness

On top of the correlations reported above, there was a moderate correlation between NASA Effort and MFI (Tau=0.306, p=0.018, BF_10_=3.542) and between NASA Effort and KSS (Tau=0.341, p=0.011, BF_10_=6.798). Finally, as classically reported in the literature, there was no correlation between the change in MFI and the change in performance (1-back: Tau=−0.133, p=0.323, BF_10_=0.392; N-back: Tau=−0.070, p=0.592, BF_10_=0.271).

### 2.3. Discussion

In experiment 1, we found only a moderate effect of the Stroop task on the subjective fatigue scores. However, the MFI questionnaire was provided at the end of the whole experiment and therefore, we suspected that the results were affected strongly by the performance of the intervening N-back task. In addition, MFI correlated strongly with KSS, suggesting that the participants failed to distinguish the concept of fatigue from that of sleepiness.

Subjective effort, as assessed through the NASA-TLX questionnaire, increased following the Stroop task, and its change correlated with that of MFI, apparently confirming our hypothesis that MF increases subjective effort cost. However, our other estimate of effort, task avoidance, correlated with objective (N-back performance), rather than subjective fatigue (MFI). One issue with the task avoidance estimate was that many participants always selected the difficult task, providing us with no information about their actual estimation of effort cost. This was presumably caused by their excessive motivation for the monetary reward.

To address these multiple methodological issues, we ran a second experiment. The confound between sleepiness and fatigue was addressed by explaining in detail to each participant the difference between the 2 concepts (Borragán et al., 2017). In addition, in order to restore vigilance, we added an “arousal boost” after the Stroop and documentary-watching sessions and before the final evaluation. Also, we measured arousal by means of pupillometry. One supplementary MFI assessment was added also right after the Stroop and documentary-watching sessions. The assessment of task avoidance was improved by using virtual coins, rather than directly euros as rewards for the participants. These coins were translated into actual money at the end of the experiment but the participants were not aware of the conversion ratio. Moreover, instead of using all the combinations of difficulty and reward, we used an adaptive procedure to choose the conditions in each trial. Finally, the selection of N-back task difficulty during the training session was also less constrained, with no maximal value for the ‘N’ of the N-back.

## 3. Experiment 2

### 3.1. Materials and methods

Thirty right-handed healthy participants took part in Experiment 2 (19 F, 11 M, Age: 23.9±4.0). MFI and KSS were assessed at the beginning and at the end of both experimental days, and between the fatigue/control manipulation and effort-discounting task, following an arousal boost which consisted in running on the spot for 30 seconds. Each participant read the NASA-TLX instructions carefully prior to the experiment, in order to ensure their appropriate understanding of the different subscales.

The first phase of the training session in Experiment 2 was similar to that of Experiment 1 and comprised 10 blocks, now with 80 letters each. Participants were then engaged in 6 blocks of N-back task with mixed difficulties, similar to those experienced during the following experimental sessions. Different ratios of 1 back & N-back tasks were performed, while the difficulty level of the N-back (i.e. the value of the N) was adapted as a function of participants' individual ability. One hundred and twenty letters were presented in these blocks. Only 3 DL were now evaluated (% of N-back: DL-1 = 20%, DL-2 = 50%, DL-3 = 80%). The monetary incentive was switched from actual money to a point-based system, and the conversion rate was not disclosed in advance to the participants.

At the beginning of each experimental session, participants performed one warm-up block (DL-3). They then had to respond to 40 forced choices, out of which 10 randomly selected blocks were actually performed.

Instead of an exhaustive screening of all possible permutations between difficulty and reward, an adaptive procedure was implemented (Kontsevich and Tyler, 1999; Watson and Pelli, 1983). Participants’ choices were modeled with the same 2-parameter effort-based decision making model as described in Experiment 1. The prior probability distribution P_0_ of the weight and temperature parameters were set to a normal distribution with mean zero and standard deviation equal to 5. In each trial, this probability distribution was updated as a function of participant’s choice:

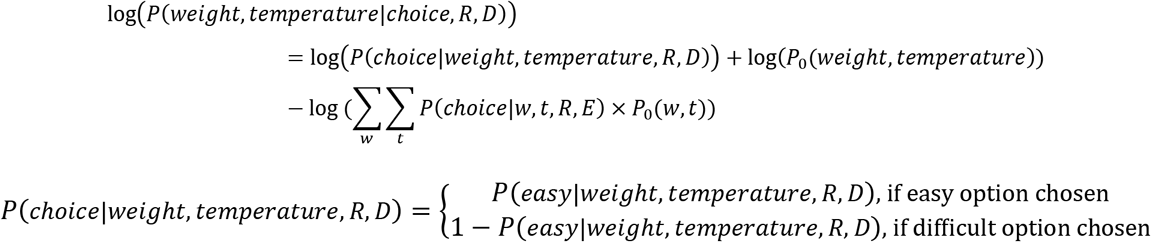

Then the expected amount of information gained in the next trial was computed for each condition, each weight and temperature values and each possible choice, and allowed us to extract a probability of choosing a given reward and difficulty condition, *P*(*R,D*), according to the following equations:

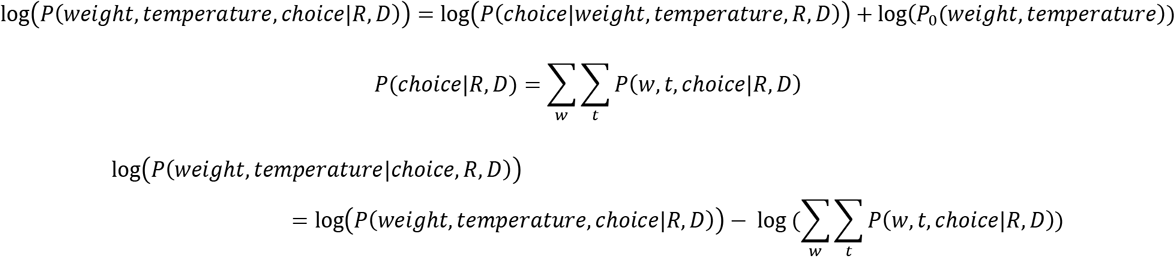

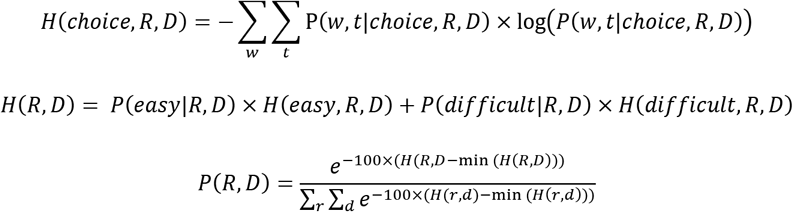

All other parameters and procedures were identical to Experiment 1.

In order to monitor participant’s level of arousal, we recorded pupil size during performance of the N-back task. Pupil size was acquired using an Eyelink 1000+ eye tracker video-based system (SR Research Ltd., Kanata, Ontario, Canada), with a sampling frequency of 500 Hz. We also evaluated maximal pupil dilation by asking the subjects to apply physical force through a handgrip on a dynamometer (Jamar^®^ Hydraulic Hand Dynamometer). This step was performed twice at the beginning and at the end of both experimental days. No pupillometry data were acquired on the training day.

### 3.2. Results

#### Stroop session increases MFI, decreases performance but leaves task avoidance unchanged

A repeated-measure ANOVA on the post-pre difference in MFI and KSS scores showed a strong effect of SESSION on KSS (see Figure 3D; F(1,29)=10.69, p=0.003, BF_10_=15.32) and a decisive effect on the MFI (F(1,29)=23.06, p<0.001, BF_10_=408.2). There was a moderate correlation between the fatigue-control changes in the two variables (Kendall's tau=0.303, p=0.026, BF_10_=3.377). Similarly to Experiment 1, there was neither SESSION effect on IMI (F(1,29)=1.965, p<0.172, BF_10_=0.657), nor correlation between IMI and MFI (Kendall's tau=0.016, p=0.900, BF_10_=0.237) or IMI and KSS (Kendall's tau=−0.015, p=0.914, BF_10_=0.237).

The NASA effort and performance scores changed as a function of the DIFFICULTY level (NASA effort: F(2,58)=26.654, p<0.001, BF_inclusion_=2.880×10^7^, NASA performance: F(2,58)=14.884, p<0.001, BF_inclusion_=10126) but not SESSION (see Figure 3F; NASA effort: F(1,29)=0.294, p=0.592, BF_inclusion_=0.203, NASA performance: F(1,29)=0.124, p=0.728, BF_inclusion_=0.171) or SESSION × DIFFICULTY interaction (NASA effort: F(2,58)=0.929, p=0.401, BF_inclusion_=0.166, NASA performance: F(2,58)=0.257, p=0.775, BF_inclusion_=0.116). There was a moderate, negative correlation between the fatigue-related changes in the two variables (Kendallɽs Tau=−0.343, p=0.007, BF_10_=7.037).

The temperature and weight parameters from the task avoidance models were also subjected to a one-way Bayesian repeated-measure ANOVA with SESSION (control vs fatigue) as factor.

There was no change in either parameters (see Figure 2, right panel and Figure 3E; temperature: F(1,29)=1.868, p=0.182, BF_10_=0.594, weight: F(1,29)=0.001, p=0.981, BF_10_=0.263) between sessions. There was no correlation between the fatigue-control changes in the two variables (Kendall's tau=0.222, p=0.092, BF_10_=0.978). The percentage of acceptance of the difficult option in Experiment 2 was 61.0±4.2% in the control and 58.4±5.4% in the fatigue session (mean±CI). None of the participants picked systematically the difficult option (all acceptance percentages were below 90%).

A RM-ANOVA on the logit-transformed accuracy scores showed, beside the difference between 1-back and N-back TASK (accuracy: F(1,29)=299.4, p<0.001, BF_10_>10^37^, RT: F(1,29)=56.587, p<0.001, BF_10_>10^15^), a moderate interaction between the TASK and the SESSION effect on accuracy (F(1,29)=7.789, p=0.009, BF_10_=4.979). This interaction was absent from the RT data (F(1,29)=0.105, p=0.748, BF_10_=0.255). The main session effect was only anecdotal (accuracy: F(1,29)=11.354, p=0.002, BF_10_=2.593, RT: F(1,29)=8.024, p=0.008, BF_10_=1.159). Looking at the accuracy in each task individually, we found a strong decrease in performance with fatigue in the 1-back task (F(1,29)=10.34, p=0.003, BF_10_=12.97) but not in the N-back task (F(1,29)=0.132, p=0.719, BF_10_=0.276). There was no correlation between the fatigue-control changes in the two variables (Kendall's tau=0.145, p=0.272, BF_10_=0.432).

#### Decrease in performance correlates with changes in task avoidance and subjective effort

We found strong evidence for a correlation between fatigue-induced changes in Model Weight and performance (see Figure 4D; N-back: Tau=−0.374, p=0.004, BF_10_=13.376; 1-back: Tau=−0.374, p=0.004, BF_10_=13.376). In contrast, Model Weight changes failed to correlate with MFI (Tau=−0.082, p=0.531, BF_10_=0.286), KSS (Tau=−0.223, p=0.110, BF_10_=0.999) or NASA score changes (see Figure 4E; effort: Tau=0.109, p=0.401, BF_10_=0.332; performance: Tau=-0.095, p=0.464, BF_10_=0.306).

There was also a positive correlation between fatigue-induced changes in performance in the N-back task and NASA performance score (see Figure 4F, blue; N-back: Tau=0.393, p=0.002, BF_10_=20.615; 1-back performance: Tau=0.264, p=0.041, BF_10_=1.784), and a moderate correlation of the performance changes with the NASA effort score (see Figure 4F, red; N-back: Tau=−0.306, p=0.018, BF_10_=3.531; 1-back: Tau=−0.149, p=0.256, BF_10_=0.450).

#### No relation between fatigue and arousal

Baseline arousal levels remained unaffected by the fatigue manipulation, as indexed by average pupil size during the MVC task (F(1,29)=0.250, p=0.621, BF_10_=0.281). The change in pupil-indexed arousal correlated moderately with the change in KSS score but not with the MFI score change (KSS: Tau=0.302, p=0.025, BF_10_=3.294; MFI: Tau=−0.005, p=0.971, BF_10_=0.235).

A 2-way Bayesian RM-ANOVA on the peak pupil response during the task showed a moderate effect of session (F(1,29)=1.912, p=0.177, BF_inclusion_=3.157), with larger pupil responses in the fatigue session. There was no DIFFICULTY (F(2,58)=1.573, p=0.216, BF_inclusion_=0.092) or interaction (F(2,58)=2.662, p=0.078, BF_inclusion_=0.245) effect. This between-session change failed to correlate with any of the fatigue measures (BF_inclusion_ between 0.2 and 0.3).

#### Subjective fatigue failed to correlate with the other fatigue or effort measures

Beside the previous correlations reported above, there was no correlation between NASA Effort and MFI (Tau=−0.012, p=0.929, BF_10_=0.236) or KSS (Tau=−0.090, p=0.504, BF_10_=0.238). Finally, there was no correlation between the change in MFI and the change in performance (1-back: Tau=−0.100, p=0.442, BF_10_=0.357; N-back: Tau=0.063, p=0.629, BF_10_=0.264).

## 4. General discussion

In the present paper, we hypothesized that cognitive fatigue would increase the cost of cognitive effort. We evaluated fatigue and effort in terms of subjective feeling, as well as objective manifestation such as task performance and reward discounting, respectively. We found that extensive involvement in a cognitively demanding task increased the feeling of fatigue and impaired task performance in the easy version of the task. Throughout the experiment, preference of the participants between task options reflected the attribution of a cost to cognitive effort (Westbrook et al., 2013). However, contrary to the hypothesis, effort cost did not increase with subjective fatigue but increased rather in proportion to fatigue-induced decrease in performance.

The cognitive mechanisms and neural substrates of effort and fatigue remain poorly understood. If subjective fatigue was a protective mechanism against forthcoming resource depletion (Gergelyfi et al., 2015; Muraven and Slessareva, 2003), or if it was the manifestation of increased engagement to maintain performance despite resource disruption (Hockey, 2013, 1997), it should also have increased task avoidance. The present findings go clearly against this hypothesis, by showing an absence of link between subjective fatigue and task avoidance. It is worth mentioning that we found a positive correlation between subjective fatigue and subjective effort in Experiment 1. However, this correlation was weak and disappeared in Experiment 2, in which we addressed multiple methodological limitations. In contrast, task avoidance correlated in both experiments with objective fatigue, i.e. the decrement in performance associated with fatigue. These findings emphasize the dissociation between subjective and objective manifestations of fatigue and effort. Indeed, in both experiments, subjective and objective assessments of fatigue and effort failed to correlate with each other.

Parts of the present findings rest on the exclusion of the alternative hypothesis. Bayesian statistics, by considering that probabilities can be attributed to hypotheses (e.g. there is a 10% chance of raining), allow us to evaluate the differences in likelihood between the null and alternative hypotheses in terms of Bayes Factor. Here, the correlation between MFI and model weight, reflecting task avoidance, resulted in both experiments in substantial evidence for the null hypothesis (Bayes Factor <0.3). This replication of the null result provides, to our view, strong evidence that MFI, as a general measure of subjective fatigue, is not associated with increased cognitive task avoidance. However, it is important to note that this finding does not demonstrate that other phenomenological aspects of fatigue, that would fail to be captured by the MFI, could not lead to increased task avoidance. Along the same line, it is possible that longer task performance, or more demanding tasks, could have led to increased task avoidance. Nevertheless, 2 hours of a task known to tax cognitive control resources, and in which difficulty was adjusted to lead to performance well below maximum, were not enough to induce increased task avoidance in the present experiment (BF below 0.3 in Exp2 and slightly above in Exp1).

We found that despite successful fatigue inducement, the participants maintained similar performance in the N-back task but showed significant decrease in 1-back performance. Our initial hypothesis was that fatigue was associated with disruption of vigilance, which typically leads to more mistakes in easy and monotonous tasks (Thiffault and Bergeron, 2003). We accounted for this possibility in Experiment 2, by adding an arousal boost manipulation to restore vigilance, by improving the dissociation between fatigue and sleepiness self-evaluation in participants, and by measuring participants’ pupil size (Sara and Bouret, 2012; Zénon et al., 2014). We found neither change of arousal between the sessions nor any correlation with the measures of fatigue, sleepiness and performance. This suggests that the decrement in 1-back performance is not caused by decreased vigilance. Therefore, understanding why fatigue disrupts specifically 1-back performance will require further investigation.

Important methodological limitations from Experiment 1 were addressed in the second experiment. After observing that effort cost was not affected by fatigue, and that many participants never selected the easy task, we hypothesized that the N-back task may not have been challenging enough and that the value of reward was too high relative to effort cost.

Therefore, in Experiment 2 we did not restrict the difficulty of the working memory task, such that some participants were able to perform a 9-back version of the task. In order to decrease the incentive value of reward, we switched from using actual money to a point-based system. Finally, instead of testing exhaustively every possible combination of task difficulty and reward, we implemented a Bayesian search procedure, in which the proposed options were adapted based on the previous responses. Despite these improvements, one remaining limitation is noteworthy. We tested young university students, who are generally strongly motivated by reward and this may have masked partly the increases in effort costs associated with fatigue.

One potential issue with the neuroeconomical evaluation procedure is that the devaluation of reward value associated with task difficulty may be influenced by performance in the task, and not only by effort. This issue deserves a careful consideration in the present study, given that we found a correlation between fatigue-induced changes in task avoidance and performance. Previous attempts at solving this issue were based on persuading participants that their reward depended on task engagement, rather than on performance (Westbrook et al., 2013). Here, we relied on the design of the task-choice procedure to avoid any influence of performance on participants’ decisions. Since block performance was computed by averaging accuracy in both the difficult and the easy tasks irrespective of their proportion in the block, a poor performance in the difficult task would have the same consequence on the reward, whatever choice the participants made. Therefore, there was no incentive for the participants to change their preference for the easy task ratio following fatigue. In fact, if participants task-choice depended on auto-evaluation of performance, decreased performance in N-back would have increased preference for low DL task conditions, whereas decreased performance in 1-back would have increased preference for large DL tasks. However, we found that performance in both the 1-back and N-back tasks correlated with task avoidance. This indicates that task avoidance was affected by fatigue-induced performance decline rather than by the integration of performance as a cost in participants’ decision. One question to address in future studies is whether the presence of the performance feedback at the end of each block is necessary to induce this relationship between performance decrements and task avoidance.

Another possible limitation of the present study is the reliance on the Stroop task to induce fatigue and N-back tasks to measure its effect. This choice was justified on the basis of several earlier studies of fatigue and effort discounting (Moeller et al., 2012; Pageaux et al., 2015; Wang et al., 2016; Westbrook et al., 2013). In addition, since we were interested in the link between the subjective feeling of fatigue - a task-independent subjective percept - and cognitive effort, there was no need to use the same task to induce and measure fatigue. Using different tasks has also the advantage of limiting the problem of boredom associated with the performance of the same monotonous task for several hours (Bench and Lench, 2013). Along the same line, it could be argued that the N-back task was not demanding enough to lead to measurable effects on effort cost. We think it is unlikely because hit rate was only 80% on average in Experiment 2 and we found clear effect of task difficulty on reward discounting.

It has been shown frequently that the feeling of fatigue occurs before any decrease of performance (Mullette-Gillman et al., 2015). Compensatory control hypothesis states that during the initial phase of cognitive fatigue, performance can be maintained at the cost of higher subjective effort (Hockey, 2013, 1997). However, the results of the present study failed to corroborate this hypothesis by showing a clear lack of relationship between subjective fatigue and effort. We believe that further computational and neuroimaging studies are required to gain a better understanding of effort cost, which would help to elucidate the intricate relationship between mental fatigue and effort.

## Acknowledgements

This work was supported by Fonds de la Recherche Scientifique (FNRS-FDP, Fondation Médicale Reine Elisabeth (FMRE), the Fondation Louvain and IdEx Bordeaux. The authors would like to acknowledge the contribution of Simon Van Hemelrijck for his support in data acquisition.

